# 3D cortical microtissue with innate microglia for studying real-time cell behavior across maturation and inflammatory response

**DOI:** 10.64898/2026.05.06.723271

**Authors:** Alexander Del Toro, Kaylen Aguilar, Angelina Clark, Alexander Bautista, Nathan Ashby, Diane Hoffman-Kim

**Affiliations:** Department of Neuroscience, Brown University, Providence, RI, USA; Carney Institute for Brain Science, Brown University, Providence, RI, USA; Institute for Biology, Engineering and Medicine, Brown University, Providence, RI, USA

**Keywords:** microglia, 3D in vitro neuroimmune model, live imaging, confocal microscopy, laser microirradiation, immune surveillance, motility, injury, inflammation, development, microtissues

## Abstract

Microglia represent the immune component of the central nervous system (CNS) that displays dynamic responses to injury and disease. Across the developing and mature CNS, microglia emerge as immunocompetent cells that continuously survey their surroundings to maintain tissue homeostasis and respond to threats. There remains a gap in 3D in vitro models that contain microglia and can provide both developmental and mature functional hallmarks. Using a 3D neural multicellular model, cortical microtissues, derived from primary rat cortical cells, we conducted live imaging to monitor microglia dynamics from early, middle, and late stage microtissue maturation. We optimized a within-micromold imaging approach that allows for live microglia imaging without removing microtissues from their culturing environment. We confirm that microglia exhibit baseline surveillance characterized by relatively stationary somas and highly dynamic cell processes that continuously extend and retract. Following proinflammatory challenges, microglia engulf lipopolysaccharide particles, accompanied by dynamic shifts in motility patterns; and rapidly respond to laser-induced tissue damage through process extension, whole-cell displacement, and local recruitment. Lastly, we show that microtissue age in culture strongly influences both baseline and directed motility profiles. Collectively, these studies demonstrate that within a 3D microenvironment, microglia exhibit pronounced changes in morphology, surveillance area, motility, and injury response across microtissue maturation. Microtissues can serve as a valuable in vitro platform for both microglia developmental studies and investigations of brain inflammation related to CNS injuries, infections, and diseases.

## Introduction

As immunocompetent cells of the central nervous system (CNS), microglia mediate key neuro-immune signals related to CNS infection, injury, and disease [1,2]. Microglia shape and maintain the developing and mature CNS through synaptic pruning and monitoring, immune surveillance, antigen presentation, chemotaxis, phagocytosis, and cytokine secretion [3–7]. Immune challenges have been shown to alter morphological, molecular, and functional microglia signatures [8–10]. Investigating microglia dynamics is vital for understanding their role in neuro-immune interactions.

Live imaging can provide a window into the diverse and dynamic repertoire of cellular behaviors corresponding to microglia states across neurodevelopment and inflammation, but it can be challenging to observe these cells in their endogenous state [11]. *In vivo* and *ex vivo* studies have established microglia as highly motile with dynamic scanning of the parenchyma for potential threats [12–15]. Live imaging in Two-dimensional (2D) culture systems have shown microglia’s role in baseline surveillance, where cellular processes and filopodia work together to maximize tissue coverage [7,16–18].

Bridging the gap between in vitro and in vivo models, 3D culture systems such as organoids and spheroids can mimic CNS features including cell composition, cell density, and electrical activity [19–21]. In some 3D in vitro neural models, exogenous microglia have been added to mature microtissues, while in others, microglia have been induced to grow in the developing culture system (for review, [22]). Current 3D models containing microglia are emerging with improved resolution for capturing real time behaviors of functions such as motility, migration, phagocytosis, and cell-cell interactions [23–30]. Despite these advances, there is a need for 3D neuro-immune models that incorporate microglia and recapitulate key functional hallmarks across developmental stages [31,32]. In addition, because of the complexity of neuro-immune interactions and especially their changes with time, there is a demand for complementary models to study microglia dynamics. A 3D neuro-immune experimental platform that can provide both developmental and mature hallmarks of microglia functionality would be of value for studying fundamental neuro-immune interactions and neuro-immune responses to disorders and injuries.

Here, we report a multicellular 3D neuro-immune model, cortical microtissues, to investigate real-time microglia behaviors across microtissue maturation, inflammation, and injury. We have used self-assembled cortical microtissues from postnatal rat cortex that show multiple CNS features including cell density, tissue stiffness, multicellular diversity, mature electrical activity, synchronous neural oscillations, myelination, and capillary-like structures. These microtissues have been used to model neural insults such as stroke, brain cancer and brain injury [33–38].

In this study, microglia in cortical microtissues were characterized from 1 to 60 days in vitro (DIV), and their morphology and motility showed hallmarks of immune surveillance behaviors that emerged across microtissue maturation. Lipopolysaccharide treatment resulted in phagocytosis by microglia and altered microglia motility dynamics in the microtissues. When exposed to laser microirradiation injuries, local microglia in microtissues were directed to injury sites. This injury responsiveness varied with the age in vitro of the microtissue. Our results suggest that cortical microtissues, with their heterogeneous composition of neural cell types, including innate microglia, reproduce key features of the CNS environment to provide a platform for studying real-time neuro-immune dynamics, that can be used for understanding microglia physiology across CNS development, injury, and neuroinflammation.

## Materials and Methods

### Rat Cortical Cell Isolation

All animal procedures were conducted according to the guidelines established by the NIH and approved by the Brown University Institutional Animal Care and Use Committee. All cortical isolation procedures followed the protocol described in Dingle et al. [33]. Briefly, male and female postnatal day 0–3 CD rat pups (Charles River) were used in all experiments; anogenital distance was used to determine sex. Cortical dissection was performed in a buffer of Hibernate A (Transnetyx HA) with 1X B27+ supplement (Invitrogen), and 0.5 mM Gluta-Max (Invitrogen). Rat cortices were cut into small pieces and digested for 30 minutes at 30 °C in 2 mg/mL papain (Transnetyx PAP) dissolved in Hibernate A without Calcium (Transnetyx HACA). After the removal of papain, tissues were triturated with a fire-polished pipette in Hibernate A buffer solution. Cell suspension was filtered through a 40 μm cell strainer and subsequently centrifuged at 150 xG for 5 min. The pellet was then resuspended in complete cortical medium + (CCM+): Neurobasal A medium (Invitrogen) with 1X B27+, 0.5 mM Gluta-Max, and 1X Penicillin-Streptomycin (Invitrogen). The suspension was then filtered, subsequently centrifuged, resuspended, and centrifuged again. Cell count and viability were determined using a trypan blue exclusion assay (Invitrogen). 2D viability control cultures were seeded in 24-well cell culture plates on wells coated with poly-D-lysine (Sigma-Aldrich) at a density of 300,000 cells/well.

### 3D Cortical Microtissue Culture

Autoclaved-sterilized powder UltraPure agarose (Invitrogen) was dissolved via heating to 2% wt/vol in phosphate-buffered saline (PBS) and poured into spheroid micromolds with 96 round pegs of 400 μm diameter (#24-96-Small, MicroTissues, Inc). After cooling, agarose gels were removed from the molds and transferred to 24-well cell culture plates. Agarose gels were equilibrated in incomplete cortical medium + (CCM+ without B27+) with 2 exchanges over the week before dissection. Gels were equilibrated in CCM + the day of dissection. For seeding, the appropriate number of cortical cells was resuspended in cortical medium at a volume of 75 μL per agarose gel. Equilibration medium was aspirated from the gels, and 75 μL of cell suspension was added to each agarose gel. Cortical cells were allowed to settle into microwells for 60 min in a 37 °C incubator with 5% *Co*_2_ with plates shaken every 20 minutes. Once cells were settled, 1 mL of CCM+ was added to each well and placed in a 37 °C incubator with a 5% *Co*_2_. Cell culture media was changed at 24 hours and every 2–3 days.

### Immunohistochemistry

Microtissues were fixed by replacing media with a 4% paraformaldehyde/8% sucrose solution overnight. The next day, the gels were washed 3 times with phosphate buffered saline (PBS) every 30 minutes. Microtissues were then stored at 4°C until staining. All of the following steps were performed on a rocking shaker at room temperature. The following antibodies were used: mouse anti-ionized calcium-binding adaptor molecule 1 (IBA1) at a concentration of 1:200 (Fujifilm 018-28523), Cy3 goat anti-Rabbit at a concentration of 1:500 (Jackson 115-165-144), and Isolectin GS-IB4 (IB4) from Griffonia simplicifolia, Alexa Fluor™ 647 Conjugate (Thermo I32450) at a concentration of 1:200. Microtissues were permeabilized and blocked with 3D Blocking Solution which consisted of 1% Triton X-100 (TX), 10% normal goat serum, and 4% bovine serum albumin in PBS for 2 hours, and subsequently incubated in primary antibodies diluted in 3D Blocking Solution overnight. Microtissues underwent two 2-hour washes with 0.2% TX in PBS (PBT), followed by one 2-hour 3D Blocking Solution wash. Microtissues were incubated with secondary antibodies in 3D Blocking Solution overnight. The Isolectin GS-IB4 from Griffonia simplicifolia, Alexa Fluor™ 647 Conjugate (1:200, Thermo) was used to label microglia and was added during secondary antibody incubation. Microtissues underwent two 2-hour PBT washes and were incubated in 1 μg/mL 4’,6-diamidino-2-phenylindole (DAPI) in PBT for 1 hour and returned to PBS before tissue clearing.

### Tissue Clearing

Before imaging fixed microtissues, samples were cleared using an adapted protocol using the Clear T2 method described in Boutin et al [34]. All of the following steps were performed on a rocking shaker at room temperature. Briefly, on the day of imaging, microtissues were placed in 10% polyethylene glycol (Sigma P2139) and 25% formamide (Sigma F9037) solution for 15 minutes. The solution was removed and replaced with 20% polyethylene glycol and 50% formamide solution for 1 hour. After 1 hour, microtissues were transferred to a glass-bottom FluoroDish (WPI FD35) for imaging.

### Fixed Imaging

Fixed microtissues were imaged using a Nikon AX-R laser scanning confocal microscope and the Nikon NIS Elements AR software. Fluorescent images were acquired from fixed and cleared microtissues using a 40X oil objective (Nikon Plan Fluor 40x Oil DIC H N2, NA 49 1.30, WD .20 mm) and laser diode (405, 488, and 532 nm) laser lines with 2x digital zoom. Image stacks were converted to TIFF format and max z-projections were reconstructed in ImageJ.

### Isolectin IB4 Dye Labeling

The Isolectin GS-IB4 (IB4) from Griffonia simplicifolia, Alexa Fluor™ 647 Conjugate was used to stain, visualize, and monitor microglia in real time [39–41]. IB4 was diluted in cell media at a final concentration of 40 μg/ml and incubated with microtissues for 1 hour. After one hour, IB4 was removed and microtissues were washed with normal cell media three times. Microtissues were then transferred to a glass-bottom FluoroDish and in an environment controlled chamber for live imaging.

### Live Imaging

Live microtissues were imaged using a Nikon AX-R laser scanning confocal microscope and the Nikon NIS Elements AR software in a 37 °C chamber with 5% *Co*_2_. Time-lapse images were acquired using the 10x air objective (Nikon Plan APO ʎD 10x OFN25 DIC N1, NA 0.45, WD 4 mm) and laser diodes (405, 488, and 532 nm) laser lines with 8 × digital zoom. For 3D live imaging, a z-stack and time-lapse images were acquired for every microtissue. We used a time acquisition interval of 2 minutes, a 3.7 micron step size, and 5 z-slices. Imaging settings were consistent between samples for all conditions within the same experiment. Image stacks were converted to TIFF format and max z-projections were reconstructed in ImageJ.

### Focal Laser Microirradiation-Induced Injury

Laser microirradiation was used to induce microtissue damage using the 405 nm laser from the Nikon AX-R laser scanning confocal microscope [42]. The 405 nm laser was set to 25% power for 16ms where each configuration occurred for 30 seconds. A single z-plane was chosen that had the largest amount of IB4+ cells. Microglia baseline was imaged from 0-5 minutes and post injury responses were imaged from 6-16 minutes.

### Lipopolysaccharide (LPS) treatment

To induce a proinflammatory challenge to microtissues, lipopolysaccharide (LPS) from Escherichia coli O111: B4 (Sigma L4391) was dissolved in cell media at a final concentration of 50 μg/ml prior to microtissue delivery.

### Lipopolysaccharide fluorescent labeling

To fluorescently label LPS particles, lipopolysaccharide (LPS) from *Escherichia coli* O111: B4 (Sigma L4391) and lipopolysaccharides Alexa Fluor™ 488 Conjugate (Thermo L23351) were combined to a 50 μg/ml and 10 μg/ml final concentration in cell media, respectively. Prior to microtissue delivery, the media with LPS and 488 conjugate solution was vortexed for 15 seconds.

### Adenosine 5′-triphosphate (ATP) treatment

To treat microtissues with Adenosine 5′-triphosphate (ATP), ATP disodium salt hydrate (Sigma) was dissolved in cell media at a final concentration of 50μM prior to microtissue delivery.

### Image Analysis

#### MotiQ

Microglia motility and morphology was quantified using the open-source plugin MotiQ available in Image J (Hansen et al.). Five to ten microglia cells were randomly selected and analyzed per microtissue. Single microglia cells were isolated using the MotiQ cropper function. Single-cell microglia images were then processed and segmented with MotiQ thresholder (selected algorithm: “minError”). Lastly, all thresholded images were analyzed using the MotiQ 3D analyzer function. Both morphological and motility parameters were used in the quantification of microglia.

#### TrackMate

Microglia displacement and velocity was analyzed using the open-source plugin TrackMate available in Image J [44,45]. The entire imaging session of laser microirradiation experiments was analyzed to assess microglia before and after laser injury. The LoG detector filter was applied for every image with a 14 µm object diameter and quality threshold of 2. The Simple LAP tracker was applied at 30 µm to the following: linking max distance, gap-closing distance, and gap-closing max frame gap. We conducted a branch hierarchy analysis within TackMate to quantify the total distance traveled and average velocity for IB4+ individual cells before and after laser microirradiation.

#### NIS Elements AR

Microglia reactivity response was measured with the Nikon NIS elements software. Coordinates from the stimulation paradigm were used as a region of interest to measure the relative fluorescence. A time series measurement was conducted for the Alexa Fluor 647 (microglia) channel across the entire image series.

#### Statistics and data visualization

Data visualizations and statistical analysis were performed using GraphPad Prism 9 (GraphPad Software LLC). All data analysis was performed using a two-tailed unpaired t-test, one-way ANOVA with post-hoc Tukey-HSD tests, or two-way ANOVA with post-hoc Sidak test.

## Results

### Microglia in cortical microtissues were monitored in real-time

Live imaging allows for studying real-time cellular dynamics and behaviors of microglia [11]. Here we optimized a within-micromold imaging pipeline that allows for monitoring microglia in arrays of microtissues without the removal from their microenvironment (Fig 1A, <u>Supp Video 1</u>). To visualize primary microglia in multicellular microtissues, we used the fluorescent dye isolectin IB4 and monitored cellular dynamics using 3D time-lapse confocal microscopy (Fig 1B). Double-staining of microtissues with immunofluorescence for microglial marker IBA1 and fluorescent isolectin IB4 confirmed the utility of IB4 for labeling microglia (Fig 1C). This setup allows for imaging multiple populations of microglia across microtissues and permits live imaging directly within the non-adhesive agarose micromold (Fig 1D-E), where cells are visualized with individual cell resolution. This 3D live imaging pipeline requires no harvesting of microtissues and allows for real-time acquisition of multiple populations of microglia in a controlled and reproducible manner.

**Figure 1.**
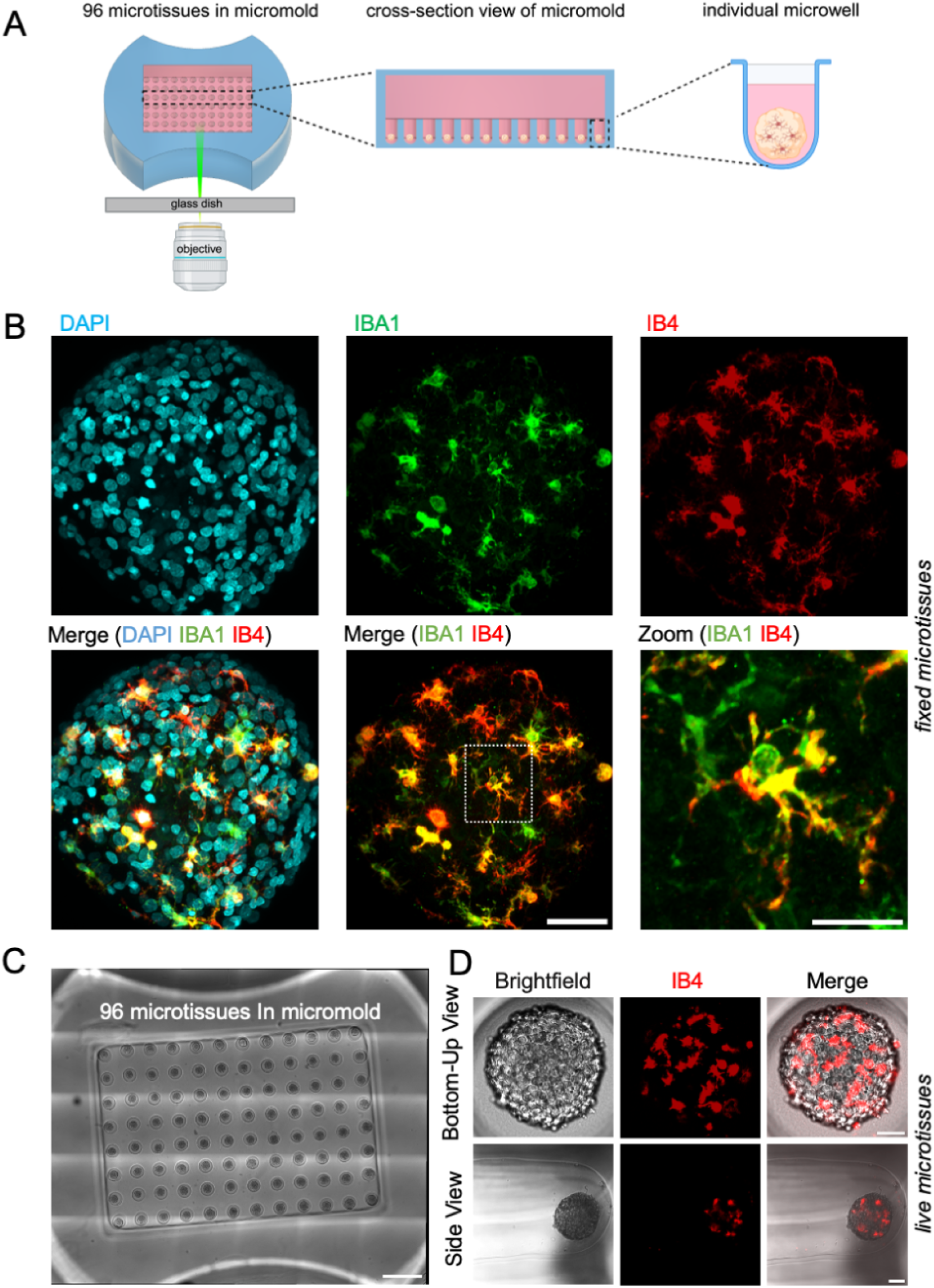
Live imaging setup to visualize and monitor microglia motility in microtissues. A Schematic image of the within-micromold live imaging setup to monitor microglia motility using confocal microscopy. B Confocal z-projections of a fixed and cleared microtissue stained with nuclear marker DAPI (cyan), microglia marker anti-IBA1 (green) antibody, and microglia marker Isolectin GS-IB4 (IB4) (red). Scale bar is 50 µm. Individual and merged fluorescent channels are shown. White box shows enlargement of an individual microglia cell co-labeled with anti-IBA1 and IB4 (yellow), shown in the high magnification image. Scale bar is 20 µm. C Confocal micrograph showing 96 live microtissues in wells of agarose micromold. Scale bar is 1000 µm. D Confocal z-projections of the Bottom-up view (top row) and Side view (bottom row) of an individual microtissue with IB4 labeled microglia. Scale bars are 50 µm.

### Microglia scanned the local microenvironment in cortical microtissues

Microglia use elaborate cellular processes to scan the extracellular space for potential threats within the CNS [12]. Under steady-state, microglia spontaneously extend and retract their processes, in baseline surveillance [12,17]. To study baseline surveillance, we used 3D time-lapse confocal imaging to monitor microglia motility dynamics in microtissues. Live microglia imaging at 15 days in vitro (DIV) within microtissues showed that microglia were generally stationary, with minimal cell displacement (<u>Supp Video 2</u>). Ramified and rounded microglia were present in microtissues and the two forms showed distinct motility dynamics (Fig 2A, <u>Supp</u> <u>Video 3</u>). We analyzed microglia using the open source software MotiQ to assess baseline dynamics [43]. Cell skeleton and size characteristics were measured by the cell volume and the number of branches for each microglia. The scanning area size was measured with the convex hull and represents the maximum microtissue volume occupied by an individual microglia cell. Dynamic characteristics such as number of extensions and retractions, connected pixels that newly emerge or disappear compared with the previous time step, correspond to a cell’s surveillance activity. Shape dynamics, the sum total of extended and retracted area volume, display the overall change of the microglia cell shape. Microglia motility in microtissues included dynamic changes in cell structure along with continuous extensions and retractions of their cellular processes (Fig 2 A-B). We quantified individual microglia dynamics (Fig 2C-D) and observed fluctuating patterns in morphology (number of branches, volume), motility (number of extensions/retractions, and shape dynamics), and scanning area size (spanned volume) (Fig 2E). Taken together, microglia in microtissues displayed a range of baseline motility behaviors that mimic a critical hallmark of microglia functionality, immune surveillance.

**Figure 2.**
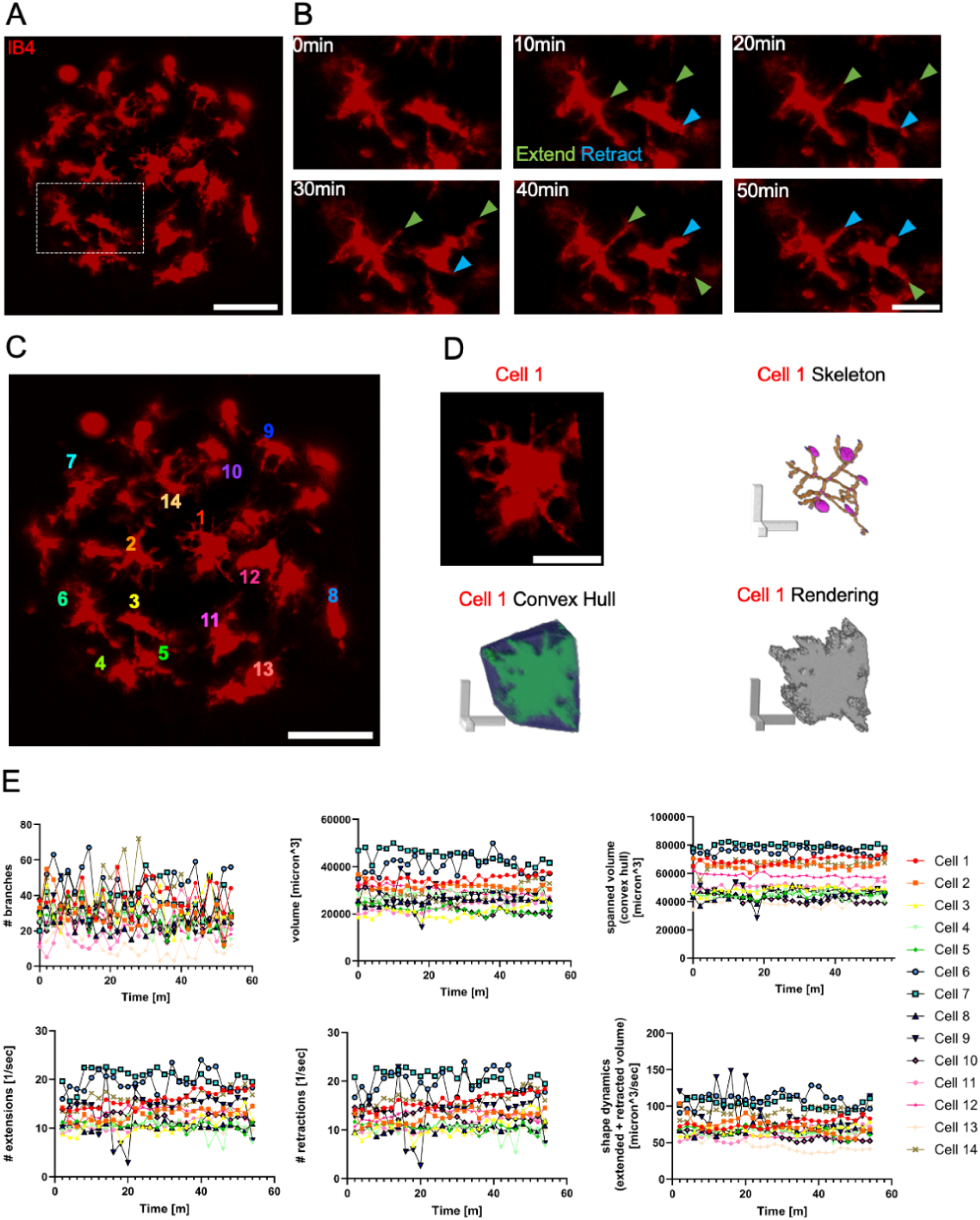
Microglia in microtissues are highly motile and scan the 3D microenvironment with dynamic cellular processes. A Confocal z-projections showing microglia labeled with IB4 in a live microtissue at 15 days in vitro (DIV). Scale bar is 50 µm. B Time series of confocal z-projections of microglia shown in white box in (A), shown at 10-minute intervals. Green arrowhead, process extension; Blue arrowhead, process retraction. Scale bar is 20 µm. C Confocal z-projection of microtissue from (A). Numbers show individual microglia for analysis in (D) and (E). Scale bar is 50 µm. D Confocal z-projection of cell 1 from (C) and 3D visualizations from MotiQ, including the 3D cell skeleton, convex hull, and 3D rendering. Scale bars are 20 µm. E Individual cell analysis of microglia in microtissue from (A-C) showing graphs of the number of branches, volume, spanned volume, number of extensions, number of retractions, and shape dynamics.

### Lipopolysaccharide treatment triggered phagocytosis and altered microglia motility dynamics in cortical microtissues

Neuroinflammation, a complex series of local immune responses in the CNS, alters microglia function [46,47]. To investigate microglia motility profiles under a pro-inflammatory state, we used a lipopolysaccharide (LPS) challenge and studied real-time dynamics of microglia in microtissues. Microglia engulfed LPS particles, and microglia clustering was observed (Fig 3A, <u>Supp Video 4</u>). Additionally, we compared microglia responses to two challenges, LPS and Adenosine 5′-triphosphate (ATP). Under baseline conditions, microglia displayed a characteristic ramified morphology (Fig. 3B, 3D). During LPS treatment, microglia exhibited reduced complexity, while during ATP treatment, microglia exhibited elongated cellular processes (Fig 3B, 3D, <u>Supp</u> <u>Video 5</u>, <u>Supp Video 6</u>). Quantification of microglia shape dynamics showed contrasting effects, with reduced dynamics during LPS treatment and increased dynamics during ATP treatment (Fig 3C, 3E). Lastly, we treated microtissues with LPS for 24 hours to study challenged microglia motility dynamics. Control microglia showed features of tiling, ramified morphology, and spontaneous scanning, whereas LPS-treated microglia showed reduced complexity, surveillance activity, and scanning area size (Fig 3F-G). Twenty-four hour LPS treatment significantly decreased microglia branches, spanned volume, number of extensions, and shape dynamics (Fig 3H-K). Taken together, microglia in microtissues could engulf foreign particles and showed sensitivity to specific treatments that trigger distinct motility profiles.

**Figure 3.**
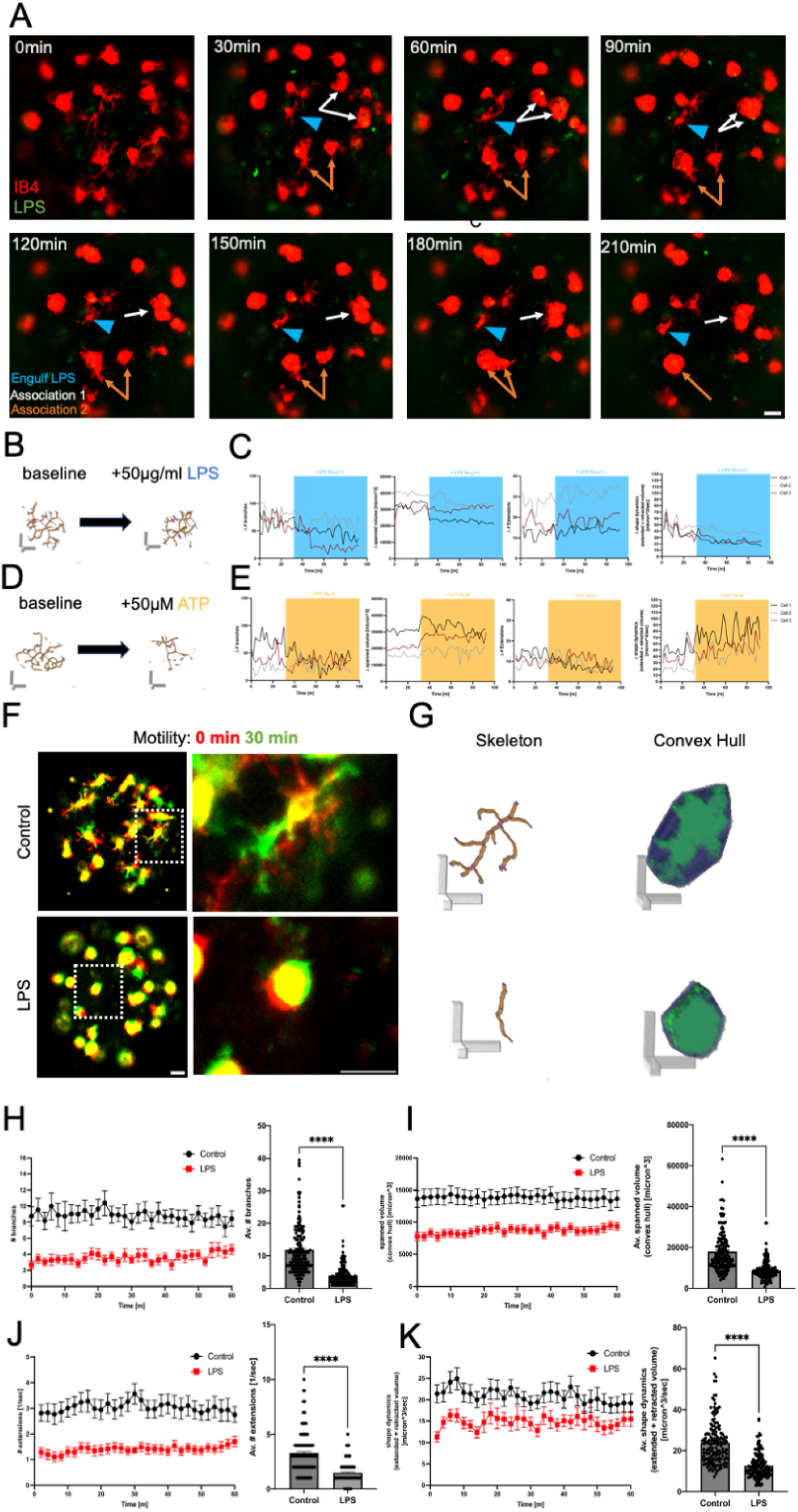
Microglia in DIV15 microtissues were phagocytic and showed morphology and motility changes following immune challenges. A Time series of confocal z-projections showing IB4 labeled microglia (red) with LPS particles (green). Blue arrowhead: microglia that engulfs LPS particle between 120-150 min; orange/white arrows: microglia that clustered near neighboring microglia. Scale bar, 50 µm. B Skeleton of representative microglia at baseline and during LPS treatment. Scale bar, 20 µm. C Graphs show individual cell traces of change in number of branches, spanned volume, number of extensions, and shape dynamics before and after LPS (blue) treatment. D Skeleton of representative microglia at baseline and during ATP treatment. Scale bar, 20 µm. E Graphs show individual cell traces for change in number of branches, spanned volume, number of extensions, and shape dynamics before and after ATP (yellow) treatment. F Representative confocal z-projections of control (top) and LPS-treated (bottom) microtissues with IB4-labeled microglia, at 0 min (red) and at 30 min (green). White boxes show individual cells that are enlarged in the right paired images. Yellow represents the overlap between time frames. Left scale bar, 50 µm; right scale bar, 20 µm. G 3D renderings from F of cell skeleton and convex hull. Scale bar, 20 µm. H-K Cell analysis of microglia in microtissues. Quantification of number of microglia branches (H), spanned volume (I), number of extensions/sec (J), and shape dynamics (K). In each graph pair, one point on the left graph represents the mean value from one point in the time-lapse imaging series. Black, control; red, LPS. (n = 30 microglia per condition). In each graph pair, one point on the right graph shows mean ± SEM, and one point represents the average value across all time points for an individual microglia (control, n = 150; LPS, n = 143). Statistical significance by two-tailed unpaired t-test (p < 0.05, p < 0.01, p < 0.001, p < 0.0001).

### Microglia surveillance emerged across cortical microtissue maturation

Microglia surveillance is a dynamic process that allows for the maintenance of tissue homeostasis, immune defense, and is critical for the developing and mature CNS [4,32,48]. Live in vivo imaging has shown unique profiles for morphology and baseline motility dynamics across the mammalian lifespan, suggesting that age influences baseline surveillance behaviors [49]. To investigate baseline microglia dynamics across microtissue maturation, we conducted 3D time-lapse imaging from 1 to 60 DIV in microtissues and assessed morphology, motility, and scanning area size (Fig 4A-B). Microglia became morphically complex across microtissue maturation with higher numbers of branches at DIV 15, 30 and 60 than DIV 1 (Fig 4C). While we saw no changes in overall cell volume (Fig 4D), the area of microglia scanning (spanned volume) was higher at DIV 30 than DIV 1 (Fig 4Ebottom). Surveillance activity, the numbers of extending and retracting events, was higher at DIV 15, 30, and 60 than at DIV 1. However, these numbers were lower at DIV 60 than at DIV 15 and 30 (Fig 4Etop-F). Shape dynamics, the sum of the total extended and retracted area, increased beyond DIV 1 with a significant decline at DIV 60 (Fig 4G). Together, these results show that baseline microglia morphology, motility, and scanning area size profiles changed based on microtissue maturity (<u>Supp Video 7</u>). Overall, the age of the microtissue can influence baseline hallmarks of microglia dynamics, which potentially correspond to developmental features seen with in vivo models.

**Figure 4.**
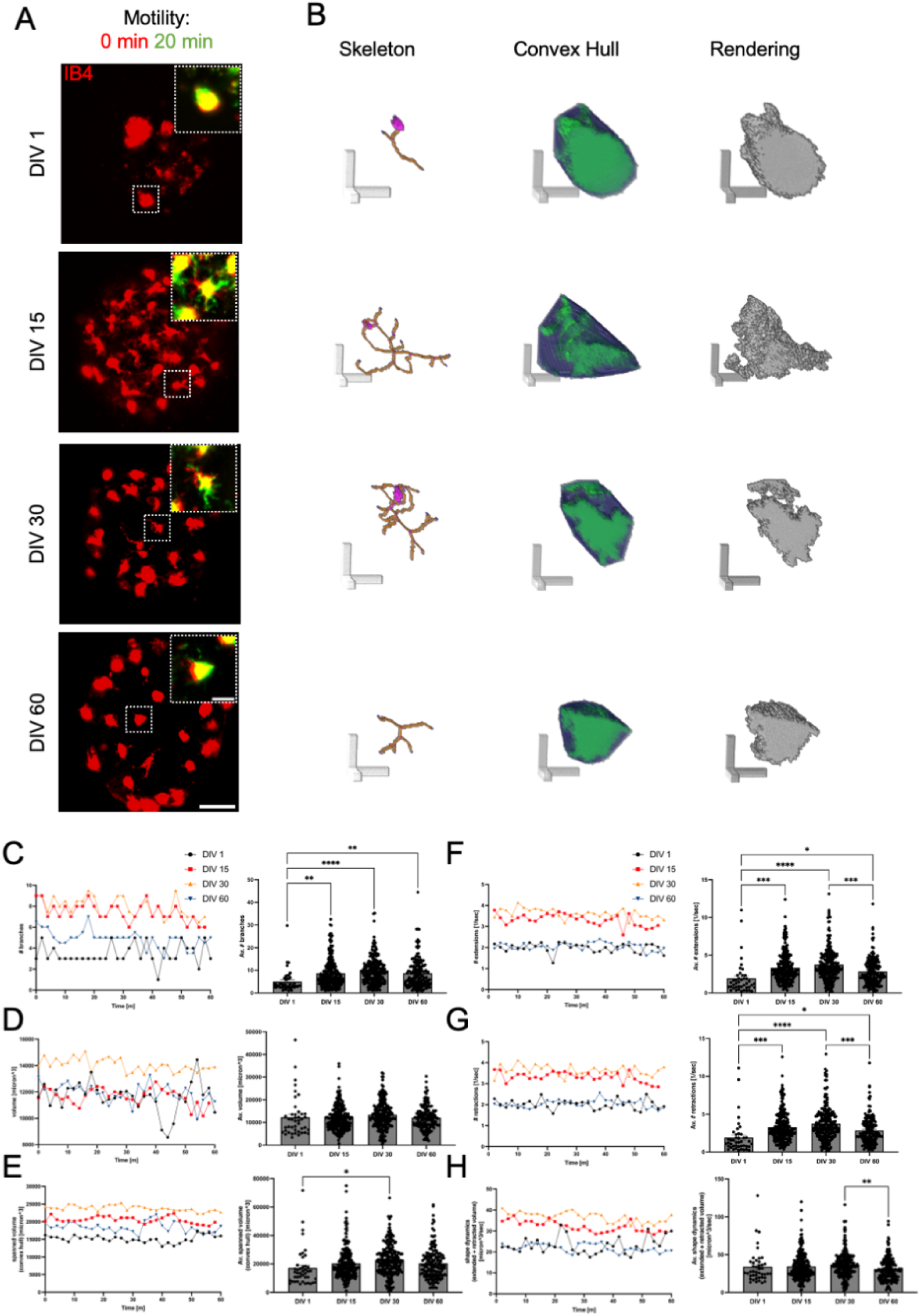
Microglia in microtissues showed distinct profiles across maturation. A Representative confocal z-projections of microtissues at 1, 15, 30 and 60 DIV with IB4-labeled microglia (red). Scale bar is 50 µm. White boxes show individual cell enlargements with 0-minute and 20-minute time frames to show cell dynamics. Yellow represents the overlap between the time frames. Scale bar is 20 µm. B Individual cell renderings from A for each DIV showing the cell skeleton, convex hull, and rendering. Scale bars are 20 µm. C-G Cell analysis of microglia in microtissues. Quantification of number of microglia branches (C), volume (D), spanned volume (E), number of extensions/sec (F), number of retractions/sec (G), and shape dynamics (H). In each graph pair, one point on the left graph represents the median value at one time point of the time-lapse imaging series. Black, DIV1; red, DIV15; orange, DIV30; blue, DIV60. *nDIV*1 = 15, *nDIV*15 = 50, *nDIV*30 = 50, *nDIV*60 = 50. In each graph pair, one point on the right graph shows mean + SEM, and one point represents the average value across all time points for an individual microglia. Statistical significance, one−way ANOVA with post−hoc Tukey test (∗p < 0.05, ∗∗p < 0.01, ∗∗∗p < 0.001, ∗∗∗∗p < 0.0001). *nDIV*1 = 43, *nDIV*15 = 230, *nDIV*30 = 228, *nDIV*60 = 166.

### Local microglia were directed to injury sites following laser microirradiation in cortical microtissues

Microglia are equipped with molecular machinery to sense local chemical changes within their microenvironment [50]. Upon neural injury, microglia can shift from baseline to a directed surveillance state, also referred to as chemotaxis, which serves as a protective response to acute neural tissue damage [13–15]. We utilized a laser microirradiation setup to study microglia motility following microtissue damage [42]. We studied microglia recruitment patterns to laser injury sites by analyzing the presence of fluorescent microglia in regions corresponding to microtissue damage. Following a 30 second laser microirradiation treatment in DIV 15 microtissues, microglia that were near the injury site exhibited directed motility toward the site (Fig 5A-B). Quantification of this motility showed that directed microglia had higher velocities than undirected microglia (Fig 5C-F). In short, local microglia can rapidly respond within minutes in a directed motility fashion in microtissues following laser-induced tissue damage.

**Figure 5.**
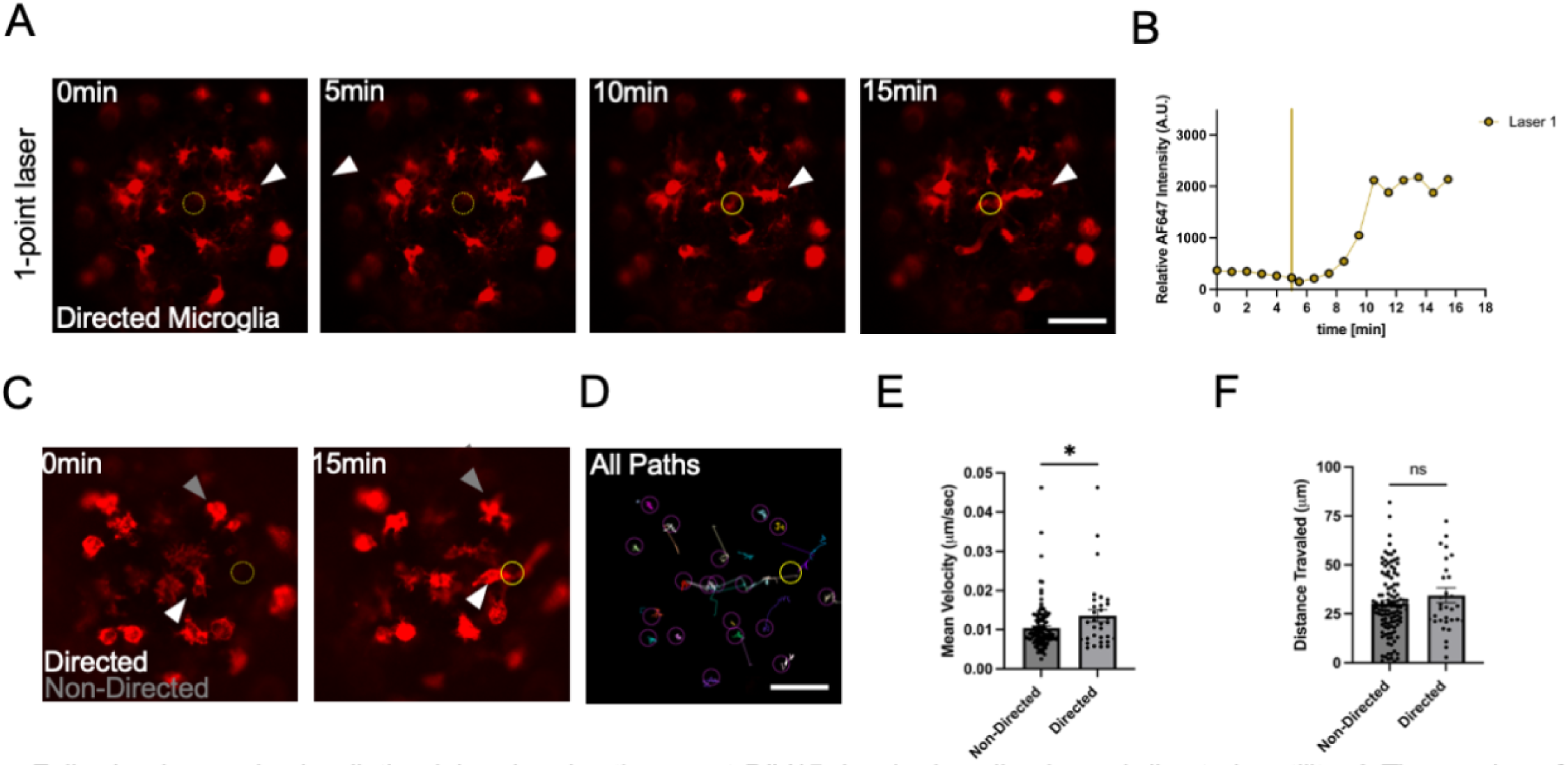
Following laser microirradiation injury in microtissues at DIV15, local microglia showed directed motility. A Time series of confocal micrographs showing IB4 labeled microglia (red). Dashed yellow circle (0 and 5 min) marks the region of upcoming laser injury, solid yellow circle (10 and 15 min) marks confirmed injury site. White arrowhead shows an individual microglia responding to laser injury. Scale bar is 50 µm. B Graph of the relative fluorescence intensity measured at the laser injury site. Yellow vertical line corresponds to laser injury pulse. C Confocal micrograph from the time series showing IB4 labeled microglia (red) pre injury (0 min) and post injury (15 min). Dashed yellow circle (0 min) marks the region of upcoming laser injury, solid yellow circle (15 min) marks confirmed injury site. Grey arrowhead shows a non-responsive microglia cell to laser injury, and white arrowhead shows a responsive microglia cell to the laser injury site. D TrackMate object pathway image showing all microglia (purple circles) and all microglia pathways (colored lines) across the time-lapse series from (C). Scale bar is 50 µm. Yellow circle is the area of injury site. E-F Quantification of mean velocity and distance traveled for directed and non-directed microglia. All points represent values for an individual microglia from 6 microtissues. Data are reported as Mean ± SEM. Statistical significance was determined by two-tailed unpaired t-test (*p<0.05). Sample sizes, N of directed microglia = 32, N of non-directed microglia = 117.

### Local microglia showed differential responses to multiple laser injuries across microtissue maturation

Recent work in animal rodent models has shown unique microglia response profiles to CNS parenchyma or vascular damage via laser injury across the mammalian lifespan, suggesting age influences patterns of directed surveillance following tissue damage [4,49]. We modified our laser injury setup with a two-point laser microirradiation injury paradigm, in which the one-point laser injury described in Figure 5 was followed immediately by another focal laser microirradiation injury at a different location in the microtissue. In DIV 15 microtissues, microglia near each injury site showed directed motility toward their respective local injury sites (Fig 6A-B). We compared the two-point laser injury responses of microglia in microtissues at different stages of in vitro maturity, and observed distinct response profiles (Fig 6C-D). The overall trends showed that microglia in microtissues at DIV 15 and 30 were significantly more responsive to injury than microglia in microtissues at DIV 2, 7, and 60. Slight differences could be seen between the responses to lasers 1 and 2 (Figure 6E). In particular, following laser 1, responsiveness emerged at the 10 minute time point, when microglia in DIV 30 microtissues were significantly more responsive than those at DIV 2 and 60 (Figure 6E). Following laser 2, responsiveness emerged at the 9 minute time point, when microglia in DIV 15 microtissues were significantly more responsive than both those at DIV 7 (Figure 6E) and those at DIV 60 (Figure 6E); and microglia in DIV 30 microtissues were significantly more responsive than those in DIV 7 microtissues (Figure 6E). In summary, the age of the microtissue shapes microglia response profiles to laser-induced tissue damage.

**Figure 6.**
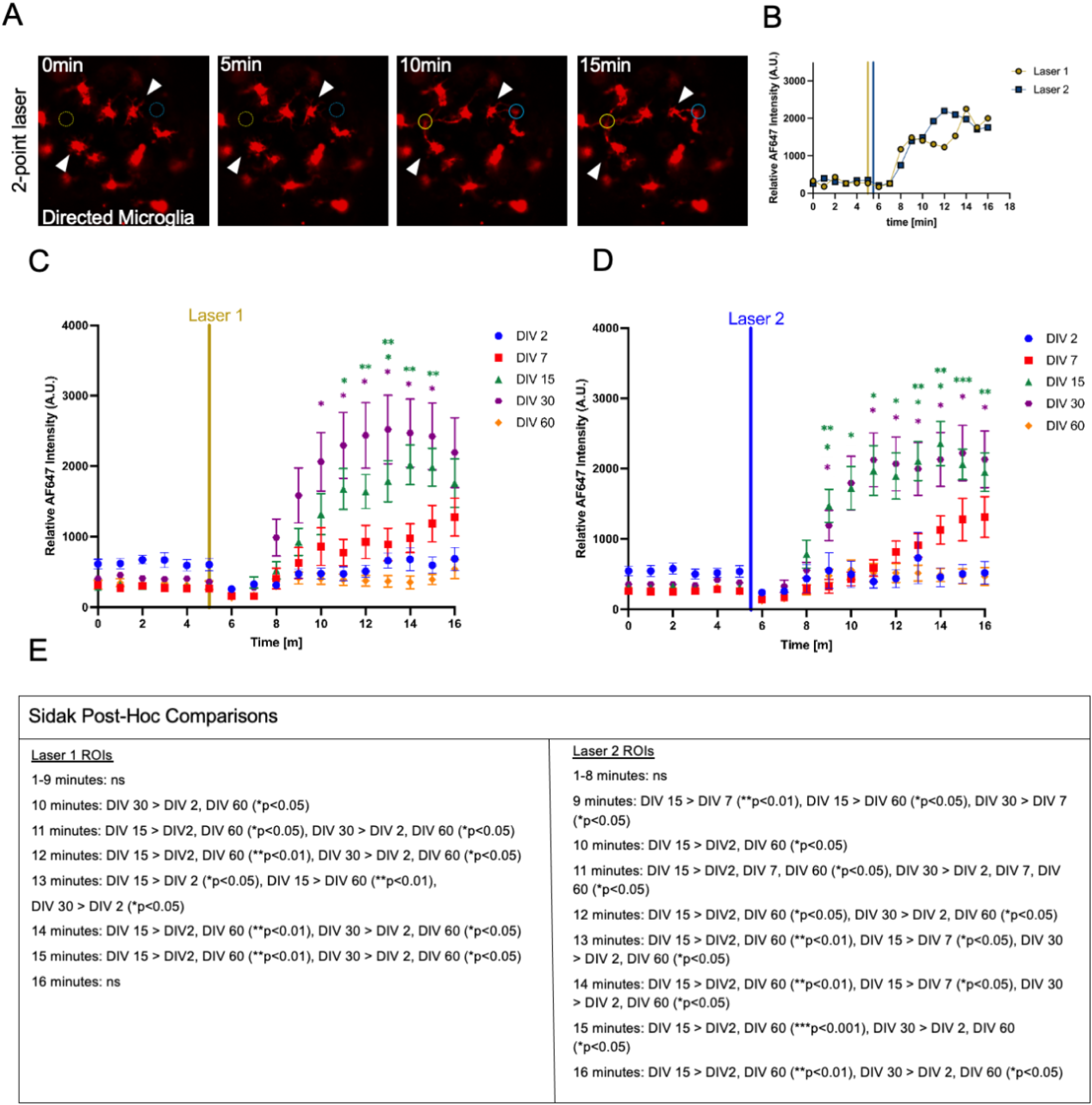
Following two-point laser injuries in microtissues, local microglia showed differential responses across microtissue maturation. A Time series of confocal micrographs showing IB4 labeled microglia (red). Dashed circles (0 and 5 min) mark regions of upcoming laser injury, while solid circles (10 and 15 min) mark confirmed injury sites. Laser sites are color-coded: yellow (laser 1), blue (laser 2). White arrowheads show representative microglia cells responding to each laser injury site. Scale bar is 50 µm. B Graph of the relative fluorescence intensity measured at each laser injury site. Vertical lines correspond to laser injury pulses. Yellow, laser 1, blue, laser 2. C-D Graph of the relative fluorescence intensities over time measured at laser site 1 (C) and at laser site 2 (D), following a two-point laser injury, for microtissues at DIV 2, 7,15, 30, and 60. All points represent averaged values from individual microtissues. Data are reported as Mean + SEM. Statistical significance was determined using a two-way ANOVA with post-hoc Sidak test. Sample sizes: N for DIV2 = 9, N for DIV7 = 10, N for DIV15 = 10, N for DIV30 = 10, N for DIV60 = 11. E Table summarizing Sidak post-hoc pairwise comparisons across time for each laser ROI. Significant differences between DIV groups are indicated with p-values; ns denotes non-significant comparisons.

## Discussion

Microglia play critical roles during neurodevelopment, for example, in neuronal maturation, neuronal circuit assembly and synaptic refinement, and during neuro-inflammatory responses to disease and injury [3,46,46,51–53]. Both developmental disorders involving microglia as well as chronic inflammation mediated by microglia have not yet achieved therapies, so experimental models that augment the current pipeline toward success are needed. In this study, we utilize a 3D neural microtissue, self-assembled from postnatal rat cortex, that contains innate microglia. This microtissue presents brain-like cell density, multicellularity, tissue stiffness, electrical activity, neuronal oscillatory behavior, extracellular matrix, and specialized structures including myelin and capillary-like networks [33–35].

We confirmed that the cortical microtissue contained microglia with immune response capabilities by treating microglia with LPS, which resulted in phagocytosis of LPS particles, reduced microglia morphological complexity, and altered motility profiles. To characterize the microglia component of the cortical microtissue during in vitro maturation, we assessed baseline microglia morphology, motility, and scanning area between 1 and 60 days in vitro (DIV), and found that microglia showed hallmarks of immune surveillance that emerged across microtissue maturation. We then modified a laser microirradiation setup to study microglia response patterns to multi-point laser stimulation in microtissues. Microglia displayed local patterns of directed surveillance to tissue damage via laser injury, where cells extended cellular processes or migrated to injury sites. Examination of this phenomenon across microtissue maturation revealed a stronger directed surveillance response from microglia in mature (DIV 15 and 30) microtissues, with weaker responses from less mature (DIV 2 and 7) or more mature (DIV 60) microtissues.

Real time imaging can illuminate the complex cell behaviors of microglia across development, surveillance, and immune response. Live imaging of microglia in 3D at individual cell resolution can pose unique optical challenges including light scattering, biomaterial penetrance, and dimensional positioning of cells [54]. Following the development of 3D neural culture systems, methods for monitoring microglia have been complicated by factors such as culture method (e.g., biomaterial, microfluidic, or scaffold-free system), and imaging system (e.g., widefield, confocal, or multiphoton microscopy) [55,56]. In the present study the cortical microtissues contained fluorescently labeled microglia in a dense multicellular environment, floating in cell culture media in a transparent agarose micromold. The imaging pipeline provided acquisition of a z-stack of images every two minutes, allowing us to image microglia in multiple microtissues with minimal photobleaching.

Before live imaging studies, microglia were thought to be resting cells waiting to be called into action. The microglia in the cortical microtissues in these experiments displayed both of the main modes of microglia surveillance: baseline and directed [12,13], which are thought to be functionally and mechanistically distinct modes [17,57] mediated by actin and tubulin cytoskeletal dynamics [58,59]. Microglia motility in microtissues was altered in the presence of a bacterial endotoxin immune challenge, and dynamic shifts in microglia motility and cell structure resulted when microtissues were treated with exogenous ATP. These results, coupled with the demonstration of microglia directed response to laser injury, suggest that microglia in cortical microtissues mimic multiple in vivo behaviors critical for immune response. It will be interesting in future studies to investigate the mechanisms underlying these distinct behaviors.

Our results show that microglia baseline surveillance was sustained across early, middle, and late stages of microtissue maturation. Motility and surveillance patterns were shaped by the age of the microtissue system, which corresponded to differential microglia motility profiles. This suggests that microglia in microtissues can show a developing phenotype, a mature phenotype, and may adopt an aging phenotype with time in culture. These results give rise to the possibility that developmental features of microtissue maturation may be preserved in a biomimetic 3D microenvironment [32,49]. What is driving these differential motility patterns is of particular area of interest for understanding microglia physiology in the context of aging or senescence-like states. Of note, the abilities of microglia to shift to a directed surveillance state following a laser injury in microtissues varied with microtissue age in culture. Microglia in mature microtissues showed directed response capabilities, while microglia in developing and older microtissues did not. The mechanisms driving baseline and directed surveillance at early and late stages of microglia maturity may provide insights into therapeutic strategies that target motility dynamics as a way to promote efficient responses in aged, injured, or diseased contexts.

## Conclusion

We have described a multicellular 3D neuroimmune microtissue model, derived from primary rodent neural cells, in which microglia innately grow, develop, and mature into surveillant, immunocompetent cells. Using 3D confocal microscopy and a within-micromold imaging approach, we optimized a platform to monitor and study microglia spatiotemporal dynamics. Each micromold contains 96 microtissues and the setup allows for live imaging. Microglia in microtissues displayed key functional hallmarks that varied with microtissue in vitro maturation age. This study demonstrates that microtissues can be used to explore the spatiotemporal dynamics of microglia, immune surveillance, and surveillant-state changes across immune challenges.

## Abbreviations

CCM+: Complete cortical medium +
PBS: Phosphate buffered saline
DAPI: 4’,6-diamidino-2-phenylindole
IB4: Isolectin GS-IB4
TX: Triton X
PBT: 0.2% Triton X in PBS
CNS: Central nervous system
2D: Two-dimensional
3D: Three-dimensional
DIV: Days in vitro
LPS: Lipopolysaccharide
ATP: Adenosine 5′-triphosphate
IBA1: Ionized calcium-binding adaptor molecule 1

## Availability of data and materials

The data used to generate all figures and videos will be held in the public repository Open Science Framework once the manuscript is accepted and published in a journal.

## Acknowledgements

We would like to thank Francesca Vecchio for providing the CAD images of the agarose micromold.

## Contributions

Conceptualization, ADT and DHK.; methodology, ADT, and DHK.; imaging, visualization, and analysis ADT, KA, AB, AC, NA.; manuscript design and organization, DHK.; writing, ADT and DHK; supervision, funding acquisition, and review and editing, DHK

## Ethics declarations

All animal procedures were conducted by the guidelines established by the NIH and approved by Brown University’s Institutional Animal Care and Use Committee [IACUC #23-07-0005].

## Consent for publication

Not applicable.

## Additional information

Publisher’s Note

## Supplementary Information

Supplementary Video 1: Video of the within micromold imaging setup to monitor real time microglia dynamics with confocal microscopy.

Supplementary Video 2: Video of IB4-labeled microglia (red) within the microtissue (brightfield) scanning the environment at 15 days in vitro.

Supplementary Video 3: Video of rounded and ramified IB4-labeled microglia (red) scanning the microtissue environment at 15 days in vitro.

Supplementary Video 4: Video of dynamics between IB4-labeled microglia (red) and LPS particles (green) in microtissues.

Supplementary Video 5: Video of dynamics before and after LPS treatment with IB4-labeled microglia (red) in microtissues.

Supplementary Video 6: Video of dynamics before and after ATP treatment with IB4-labeled microglia (red) in microtissues.

Supplementary Video 7: Video of individual IB4-labeled microglia (red) showing motility dynamics at 1,5,10,15,20,25,30, and 60 days in vitro.

## References

1. Prinz M, Jung S, Priller J. Microglia Biology: One Century of Evolving Concepts. Cell. 2019;179:292–311. 10.1016/j.cell.2019.08.053

2. Tremblay M-È, Stevens B, Sierra A, Wake H, Bessis A, Nimmerjahn A. The Role of Microglia in the Healthy Brain: Figure 1. J Neurosci. 2011;31:16064–9. 10.1523/JNEUROSCI.4158-11.2011

3. Schafer DP, Lehrman EK, Kautzman AG, Koyama R, Mardinly AR, Yamasaki R, et al. Microglia Sculpt Postnatal Neural Circuits in an Activity and Complement-Dependent Manner. Neuron. 2012;74:691–705. 10.1016/j.neuron.2012.03.026

4. Petry P, Oschwald A, Kierdorf K. Microglial tissue surveillance: The never-resting gardener in the developing and adult CNS. Eur J Immunol. 2023;53:2250232. 10.1002/eji.202250232

5. Byram SC, Carson MJ, DeBoy CA, Serpe CJ, Sanders VM, Jones KJ. CD4-Positive T Cell-Mediated Neuroprotection Requires Dual Compartment Antigen Presentation. J Neurosci. 2004;24:4333–9. 10.1523/JNEUROSCI.5276-03.2004

6. Hetier E, Ayala J, Denèfle P, Bousseau A, Rouget P, Mallat M, et al. Brain macrophages synthesize interleukin-1 and interleukin-1 mRNAs in vitro. J Neurosci Res. 1988;21:391–7. 10.1002/jnr.490210230

7. Booth PL, Eric Thomas W. Dynamic features of cells expressing macrophage properties in tissue cultures of dissociated cerebral cortex from the rat. Cell Tissue Res. 1991;266:541–51. 10.1007/BF00318596

8. Paolicelli RC, Ferretti MT. Function and Dysfunction of Microglia during Brain Development: Consequences for Synapses and Neural Circuits. Front Synaptic Neurosci [Internet]. 2017 [cited 2023 Apr 26];9. https://www.frontiersin.org/articles/10.3389/fnsyn.2017.00009. Accessed 26 Apr 2023

9. Benarroch EE. Microglia: Multiple roles in surveillance, circuit shaping, and response to injury. Neurology. 2013;81:1079–88. 10.1212/WNL.0b013e3182a4a577

10. Paolicelli RC, Sierra A, Stevens B, Tremblay M-E, Aguzzi A, Ajami B, et al. Microglia states and nomenclature: A field at its crossroads. Neuron. 2022;110:3458–83. 10.1016/j.neuron.2022.10.020

11. Andoh M, Koyama R. Assessing Microglial Dynamics by Live Imaging. Front Immunol [Internet]. 2021 [cited 2024 Jan 9];12. https://www.frontiersin.org/articles/10.3389/fimmu.2021.617564. Accessed 9 Jan 2024

12. Nimmerjahn A, Kirchhoff F, Helmchen F. Resting Microglial Cells Are Highly Dynamic Surveillants of Brain Parenchyma in Vivo. Science. 2005;308:1314–8. 10.1126/science.1110647

13. Davalos D, Grutzendler J, Yang G, Kim JV, Zuo Y, Jung S, et al. ATP mediates rapid microglial response to local brain injury in vivo. Nat Neurosci. 2005;8:752–8. 10.1038/nn1472

14. Hines DJ, Hines RM, Mulligan SJ, Macvicar BA. Microglia processes block the spread of damage in the brain and require functional chloride channels. Glia. 2009;57:1610–8. 10.1002/glia.20874

15. Haynes SE, Hollopeter G, Yang G, Kurpius D, Dailey ME, Gan W-B, et al. The P2Y12 receptor regulates microglial activation by extracellular nucleotides. Nat Neurosci. 2006;9:1512–9. 10.1038/nn1805

16. Bernier L-P, Bohlen CJ, York EM, Choi HB, Kamyabi A, Dissing-Olesen L, et al. Nanoscale Surveillance of the Brain by Microglia via cAMP-Regulated Filopodia. Cell Rep. 2019;27:2895–2908.e4. 10.1016/j.celrep.2019.05.010

17. Madry C, Kyrargyri V, Arancibia-Cárcamo IL, Jolivet R, Kohsaka S, Bryan RM, et al. Microglial Ramification, Surveillance, and Interleukin-1β Release Are Regulated by the Two-Pore Domain K+ Channel THIK-1. Neuron. 2018;97:299–312.e6. 10.1016/j.neuron.2017.12.002

18. Ward SA, Ransom PA, Booth PL, Thomas WE. Characterization of ramified microglia in tissue culture: Pinocytosis and motility. J Neurosci Res. 1991;29:13–28. 10.1002/jnr.490290103

19. Marton RM, Pașca SP. Organoid and Assembloid Technologies for Investigating Cellular Crosstalk in Human Brain Development and Disease. Trends Cell Biol. 2020;30:133–43. 10.1016/j.tcb.2019.11.004

20. Lancaster MA, Knoblich JA. Generation of cerebral organoids from human pluripotent stem cells. Nat Protoc. 2014;9:2329–40. 10.1038/nprot.2014.158

21. Pașca SP, Arlotta P, Bateup HS, Camp JG, Cappello S, Gage FH, et al. A nomenclature consensus for nervous system organoids and assembloids. Nature. 2022;609:907–10. 10.1038/s41586-022-05219-6

22. Zhang W, Jiang J, Xu Z, Yan H, Tang B, Liu C, et al. Microglia-containing human brain organoids for the study of brain development and pathology. Mol Psychiatry. 2023;28:96–107. 10.1038/s41380-022-01892-1

23. Popova G, Soliman SS, Kim CN, Keefe MG, Hennick KM, Jain S, et al. Human microglia states are conserved across experimental models and regulate neural stem cell responses in chimeric organoids. Cell Stem Cell. 2021;28:2153–2166.e6. 10.1016/j.stem.2021.08.015

24. Park DS, Kozaki T, Tiwari SK, Moreira M, Khalilnezhad A, Torta F, et al. iPS-cell-derived microglia promote brain organoid maturation via cholesterol transfer. Nature. 2023;623:397–405. 10.1038/s41586-023-06713-1

25. Sarnow K, Majercak E, Qurbonov Q, Cruzeiro GAV, Jeong D, Haque IA, et al. Neuroimmune-competent human brain organoid model of diffuse midline glioma. Neuro-Oncol. 2025;27:369–82. 10.1093/neuonc/noae245

26. Wu J, Chen X, Zhang J, Wettschurack K, Robinson M, Li W, et al. Human microglia in brain assembloids display region-specific diversity and respond to hyperexcitable neurons carrying *SCN2A* mutation. Sci Adv. 2026;12:eady2977. 10.1126/sciadv.ady2977

27. Panagiotakopoulou V, Welzer M, Ormaechea OR, Werner D, Erlebach L, Bühler A, et al. Chimeric human organoid and mouse brain slice co-cultures to study microglial function. Cell Rep. 2025;44:116656. 10.1016/j.celrep.2025.116656

28. Cakir B, Tanaka Y, Kiral FR, Xiang Y, Dagliyan O, Wang J, et al. Expression of the transcription factor PU.1 induces the generation of microglia-like cells in human cortical organoids. Nat Commun. 2022;13:430. 10.1038/s41467-022-28043-y

29. Schafer ST, Mansour AA, Schlachetzki JCM, Pena M, Ghassemzadeh S, Mitchell L, et al. An in vivo neuroimmune organoid model to study human microglia phenotypes. Cell. 2023;186:2111–2126.e20. 10.1016/j.cell.2023.04.022

30. Wendt S, Lin AJ, Ebert SN, Brennan DJ, Cai W, Bai Y, et al. A 3D human iPSC-derived multi-cell type neurosphere system to model cellular responses to chronic amyloidosis. J Neuroinflammation. 2025;22:119. 10.1186/s12974-025-03433-3

31. Cuní-López C, Stewart R, White AR, Quek H. 3D in vitro modelling of human patient microglia: A focus on clinical translation and drug development in neurodegenerative diseases. J Neuroimmunol. 2023;375:578017. 10.1016/j.jneuroim.2023.578017

32. Cuesta-Puente X, Gonzalez-Dominguez M, Pereira-Iglesias M, Perez-Arriazu N, Villegas-Zafra P, Ramos-Gonzalez P, et al. Building Immunocompetent Cerebral Organoids From a Developmental Perspective. Glia. 2025;73:2154–66. 10.1002/glia.70062

33. Dingle Y-TL, Boutin ME, Chirila AM, Livi LL, Labriola NR, Jakubek LM, et al. Three-Dimensional Neural Spheroid Culture: An *In Vitro* Model for Cortical Studies. Tissue Eng Part C Methods. 2015;21:1274–83. 10.1089/ten.tec.2015.0135

34. Boutin ME, Kramer LL, Livi LL, Brown T, Moore C, Hoffman-Kim D. A three-dimensional neural spheroid model for capillary-like network formation. J Neurosci Methods. 2018;299:55–63. 10.1016/j.jneumeth.2017.01.014

35. Sevetson JL, Theyel B, Hoffman-Kim D. Cortical spheroids display oscillatory network dynamics. Lab Chip. 2021;21:4586–95. 10.1039/D1LC00737H

36. McLaughlin RM, Top I, Laguna A, Hernandez C, Katz H, Livi LL, et al. Cortical Spheroid Model for Studying the Effects of Ischemic Brain Injury. Vitro Models. 2023;2:25–41. 10.1007/s44164-023-00046-z

37. Gonzalez-Cruz RD, Wan Y, Calvao D, Burgess AZ, Renken WK, Vecchio F, et al. Cortical spheroids show strain-dependent cell viability loss and neurite disruption following sustained compression injury [Internet]. Bioengineering; 2023 Nov. 10.1101/2023.11.15.567286

38. Calvao DJ, Vecchio F, Zein-Sabatto A, Ocitti M, Vidomlanski R, Schmidt A, et al. A brain cancer microtissue model for studying tumor cell and neural cell interactions. Sci Rep. 2025;15:36021. 10.1038/s41598-025-19982-9

39. Kaur C, Ling EA. Study of the Transformation of Amoeboid Microglial Cells into Microglia Labelled with the Isolectin Griffonia simplicifolia in Postnatal Rats. Cells Tissues Organs. 1991;142:118–25. 10.1159/000147175

40. Ernst C, Christie BR. Isolectin-IB4 as a vascular stain for the study of adult neurogenesis. J Neurosci Methods. 2006;150:138–42. 10.1016/j.jneumeth.2005.06.018

41. Streit WJ. An improved staining method for rat microglial cells using the lectin from Griffonia simplicifolia (GSA I-B4). J Histochem Cytochem. 1990;38:1683–6. 10.1177/38.11.2212623

42. Levone BR, Lombardi S, Barabino SML. Laser microirradiation as a tool to investigate the role of liquid-liquid phase separation in DNA damage repair. STAR Protoc. 2022;3:101146. 10.1016/j.xpro.2022.101146

43. Hansen JN, Brückner M, Pietrowski MJ, Jikeli JF, Plescher M, Beckert H, et al. MotiQ: an open-source toolbox to quantify the cell motility and morphology of microglia. Mogilner A, editor. Mol Biol Cell. 2022;33:ar99. 10.1091/mbc.E21-11-0585

44. Tinevez J-Y, Perry N, Schindelin J, Hoopes GM, Reynolds GD, Laplantine E, et al. TrackMate: An open and extensible platform for single-particle tracking. Methods. 2017;115:80–90. 10.1016/j.ymeth.2016.09.016

45. Ershov D, Phan M-S, Pylvänäinen JW, Rigaud SU, Le Blanc L, Charles-Orszag A, et al. TrackMate 7: integrating state-of-the-art segmentation algorithms into tracking pipelines. Nat Methods. 2022;19:829–32. 10.1038/s41592-022-01507-1

46. Hoogland ICM, Houbolt C, van Westerloo DJ, van Gool WA, van de Beek D. Systemic inflammation and microglial activation: systematic review of animal experiments. J Neuroinflammation. 2015;12:114. 10.1186/s12974-015-0332-6

47. Lu Y-C, Yeh W-C, Ohashi PS. LPS/TLR4 signal transduction pathway. Cytokine. 2008;42:145–51. 10.1016/j.cyto.2008.01.006

48. Eyo UB, Miner SA, Weiner JA, Dailey ME. Developmental Changes in Microglial Mobilization are Independent of Apoptosis in the Neonatal Mouse Hippocampus. Brain Behav Immun. 2016;55:49–59. 10.1016/j.bbi.2015.11.009

49. Tieu T, Cruz A-JN, Weinstein JR, Shih AY, Coelho-Santos V. Physiological and injury-induced microglial dynamics across the lifespan [Internet]. 2024 [cited 2025 Jun 24]. 10.1101/2024.10.02.615212

50. Hickman SE, Kingery ND, Ohsumi TK, Borowsky ML, Wang L, Means TK, et al. The microglial sensome revealed by direct RNA sequencing. Nat Neurosci. 2013;16:1896–905. 10.1038/nn.3554

51. Sierra A, Encinas JM, Deudero JJP, Chancey JH, Enikolopov G, Overstreet-Wadiche LS, et al. Microglia Shape Adult Hippocampal Neurogenesis through Apoptosis-Coupled Phagocytosis. Cell Stem Cell. 2010;7:483–95. 10.1016/j.stem.2010.08.014

52. Tremblay M-È, Lowery RL, Majewska AK. Microglial Interactions with Synapses Are Modulated by Visual Experience. Dalva M, editor. PLoS Biol. 2010;8:e1000527. 10.1371/journal.pbio.1000527

53. Stevens B, Allen NJ, Vazquez LE, Howell GR, Christopherson KS, Nouri N, et al. The Classical Complement Cascade Mediates CNS Synapse Elimination. Cell. 2007;131:1164–78. 10.1016/j.cell.2007.10.036

54. Aktories P, Petry P, Kierdorf K. Microglia in a Dish—Which Techniques Are on the Menu for Functional Studies? Front Cell Neurosci [Internet]. 2022 [cited 2024 Jan 10];16. https://www.frontiersin.org/articles/10.3389/fncel.2022.908315. Accessed 10 Jan 2024

55. Cakir B, Kiral FR, Park I-H. Advanced in vitro models: Microglia in action. Neuron. 2022;110:3444–57. 10.1016/j.neuron.2022.10.004

56. Sabate-Soler S, Bernini M, Schwamborn JC. Immunocompetent brain organoids—microglia enter the stage. Prog Biomed Eng. 2022;4:042002. 10.1088/2516-1091/ac8dcf

57. Madry C, Attwell D. Receptors, Ion Channels, and Signaling Mechanisms Underlying Microglial Dynamics. J Biol Chem. 2015;290:12443–50. 10.1074/jbc.R115.637157

58. Socodato R, Relvas JB. A cytoskeleton symphony: Actin and microtubules in microglia dynamics and aging. Prog Neurobiol. 2024;234:102586. 10.1016/j.pneurobio.2024.102586

59. Blanchoin L, Boujemaa-Paterski R, Sykes C, Plastino J. Actin Dynamics, Architecture, and Mechanics in Cell Motility. Physiol Rev. 2014;94:235–63. 10.1152/physrev.00018.2013

